# Quantitative and dynamic cell polarity tracking in plant cells

**DOI:** 10.1101/2020.09.12.294942

**Authors:** Yan Gong, Rachel Varnau, Eva-Sophie Wallner, Dominique C. Bergmann, Lily S. Cheung

## Abstract

Quantitative information on the spatiotemporal distribution of polarized proteins is central for understanding cell-fate determination, yet collecting sufficient data for statistical analysis is difficult to accomplish with manual measurements. Here we present POME, a semi-automated pipeline for the quantification of cell polarity, and demonstrate its application to a variety of developmental contexts. POME analysis reveals that during asymmetric cell divisions in the *Arabidopsis thaliana* stomatal lineage, polarity proteins BASL and BRXL2 are more asynchronous and less mutually dependent than previously thought. While their interaction is important to maintain their polar localization and recruit other effectors to regulate asymmetric cell divisions, BRXL2 polarization precedes that of BASL and can be initiated in BASL’s absence. Uncoupling of polarization from BASL activity is also seen in *Brachypodium distachyon*, where we find that the MAPKKK BdYDA1 is segregated and polarized following asymmetric division. Our results demonstrate that POME is a versatile tool, which by itself or combined with tissue-level studies and advanced microscopy techniques can help uncover new mechanisms of cell polarity.

## INTRODUCTION

Cell polarity is the central mechanism responsible for organizing a cell into subdomains with specialized functions. The subcellular enrichment of polarity proteins enables complex processes like cell migration, directional long-range signal transduction, localized cell growth, and asymmetric cell division (ACD) (Drubin & Nelson, 1996; Muroyama & Bergmann, 2019). Due to the sessile lifestyle of plants, cell polarity is particularly important for the proper execution of developmental and physiological programs. Plants need to adjust their development based on extrinsic cues, and the lack of cell migration puts greater emphasis on the orientation of ACDs and differential cell expansion for tissue patterning.

Although cell polarity can be manifested in the distribution of many components, including organelles, RNAs, and metabolites, proteins polarly distributed at the cortex play especially prominent roles in plants. These “polarity proteins” may be integral or peripheral membrane components, and are often encoded only in plant genomes. Their distribution defines cellular and tissue-level axes, and they direct the development of major body axes, tissue layers, and the distribution of specialized cell types within a tissue (Muroyama & Bergmann, 2019). For example, in the stomatal lineage of the *Arabidopsis* leaf epidermis, polarly localized BREAKING OF ASYMMETRY IN THE STOMATAL LINEAGE (BASL) and the BREVIS RADIX family (BRXf) are required to orient ACDs and ensure the daughters of such divisions take on different fates (Dong et al., 2009; Rowe et al., 2019).

Many other proteins exhibiting polarized distributions in plants have been identified over the past decade, but our understanding of the mechanisms that control their distribution is still in its infancy (Denninger et al., 2019; Dong et al., 2009; Houbaert et al., 2018; Marhava et al., 2018; Scacchi et al., 2009; Tan et al., 2020; van Dop et al., 2020; Yoshida et al., 2019; Zhang et al., 2020; Zourelidou et al., 2009). Quantifying the distribution of polar proteins in time and space is a necessary first step in understanding how cell polarity is formed and coordinated with neighboring or daughter cells. Comparing the polar domain size of different polarity proteins can help infer how protein complexes are assembled, while the temporal dynamics of the protein abundance along the membrane can reveal trafficking and post-translational regulatory mechanisms.

In animal systems, quantification of polarity proteins has already challenged old paradigms and revealed unforeseen phenomena. For example, PAR proteins in animal cells have been known for nearly 30 years. Yet, a recent 3D image segmentation and quantitative analysis of PAR proteins in the *Caenorhabditis elegans* embryo germline revealed that their polarity domain size does not scale with cell size, and this intrinsic property regulates the timing of cell fate transition (Hubatsch et al., 2019).

Similar quantitative analyses in plants would help identify critical players and novel mechanisms in the establishment of cell polarity. However, moststudiesthusfar have relied heavily on tedious visual inspection and remain qualitative (Dong et al., 2009; Marhava et al., 2018; Rowe et al., 2019; Scacchi et al., 2009; van Dop et al., 2020; Wisniewska et al., 2006; Yoshida et al., 2019; Zhang et al., 2016). Early efforts to define “polarity indexes” fall roughly into two categories: those that compare protein abundance between the peak of the polarity “crescent” and the opposite side of the cell, and those that measure the relative size of the crescent. Neither of these two types of polarity indexes captures the full picture of protein distribution. Polarity indexes that only compare protein abundance between poles may be insufficient to distinguish between the subtle contributions of the multitude of processes that shape the distribution (Denninger et al., 2019; Houbaert et al., 2018; Langowski et al., 2016; Marhavy et al., 2014; Zhang et al., 2015). On the other hand, polarity indexes that consider only the width of the crescent (Zhang et al., 2015), and do not compare protein abundance at the poles, are also insufficient to fully describe polarization, as we will show in this work.

Here, we present <u>Po</u>larity <u>Me</u>asurement (POME), an image analysis pipeline designed to measure polarity in plant cells from confocal fluorescence images. Our tool converts the measurement of individual cells into data sets that can be interrogated with a variety of statistical tools. We demonstrate the use of POME to systematically quantify the polarity of BASL and BRXf over time in the *Arabidopsis* stomatal lineage. Based on these detailed measurements, we modify previous models for how these proteins establish and maintain polarity in stomatal lineage cells. We also demonstrate how POME can quantify aspects of asymmetric protein distributions in different tissues and organisms, and again reveal that previous models for polarity establishment must be reconsidered. Finally, to facilitate community access and implementation of POME, we provide our pipeline and documentation in an accessible format.

## RESULTS AND DISCUSSION

### POME: A semi-automated tool to quantify cortical protein polarity

The *Arabidopsis* stomatal lineage is a powerful model for genetic analysis of asymmetric divisions, but mechanistic dissection of cell polarity has been challenging. BASL and BRXf polarity (demonstrated with BRX and BRXL2) is required for ACDs (Dong et al., 2009; Rowe et al., 2019), but how these proteins initiate and maintain polarization is not understood. The primary challenge hindering quantitative analysis in the stomatal lineage is the rare and transient nature of cells undergoing ACDs, which demands the imaging of a large number of samples to discern statistically significant differences between genotypes or conditions. Additionally, cell shapes in the stomatal lineage are irregular and variable, which can make drawing conclusions from visual inspection difficult and adds uncertainty to manual measurements.

To reduce the time required for quantification and to collect shape-agnostic measurements, we developed POME, a pipeline composed of an intuitive FIJI macro and an R script that automatically extracts cortical polarity information from user-defined cells. POME requires confocal images of a cell with two different channels. One of these channels must contain a uniform cell outline marker (e.g., integral plasma membrane reporter, FM 4-64, or propidium iodide). The macro then quantifies the fluorescence of cortical polar proteins in the second channel at different angles (Figure 1A-C). This simplification provides comparable results that are independent of cell shape. Users can specify the number of measurements (i.e., angles) along the membrane.

**Figure 1.**
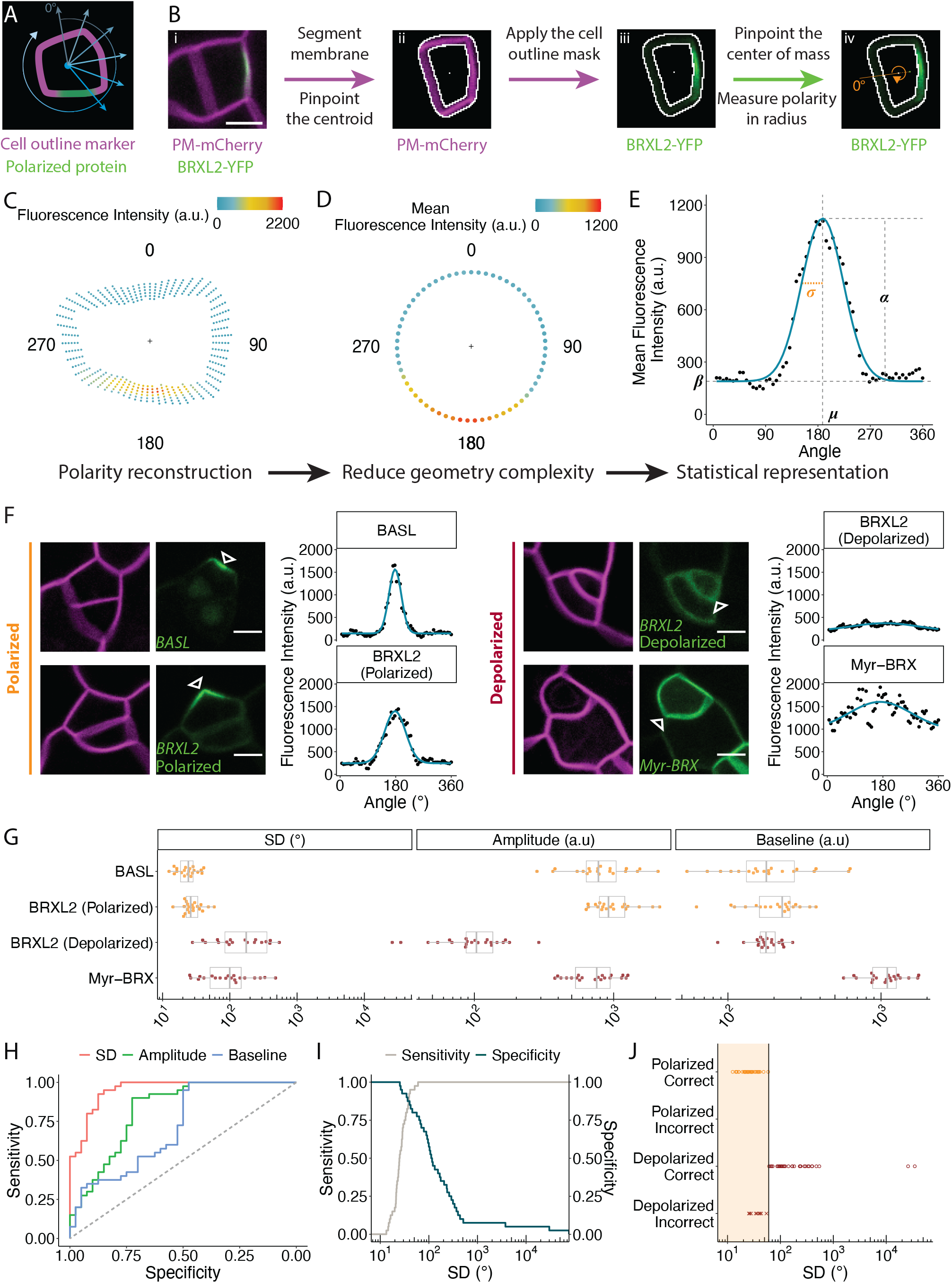
POME measures polarity in cells of the *Arabidopsis* stomatal lineage. (A) Cartoon of cell polarity analysis in POME. (B) Work flow. i) Example of a two-channel input image where one channel is a plasma membrane marker (magenta) and the second is a protein whose polarity will be quantified, in this case BRXL2-YFP (green). ii) Result of the automatic segmentation and centroid detection. iii) Application of the cell outline mask to the polarity channel. iv) Quantification of fluorescence intensity along the membrane. A line is drawn from the cell centroid (white dot in ii and iii) to the BRXL2 center of mass (green dot in iv) to define 0° and 180° angles. (C-D) Reconstruction and visualization of cell polarity from POME measurements. The cell centroid is marked with “+”. BRXL2 fluorescence intensity of each measured pixel (C) and the average BRXL2 fluorescence intensity of all the measured pixels at a given angle (D) are represented by dots colored by their fluorescence intensity. (E) Curve fitting of the average pixel intensities to a Gaussian model (blue line). The four parameters from the regression model are labelled as: standard deviation *σ*, center *μ*, amplitude *α*, and baseline intensity *β*. (F) Representative confocal images and POME measurements of polarized and depolarized markers. Magenta: *pML1::RCI2A-mCherry*; green: *pBASL::YFP-BASL* (top left), *pBRXL2::BRXL2-YFP* (bottom left and top right), *pBASL::Myr-BRX-YFP* (bottom right). Open white triangles indicate the centers of the polarity domain suggested by POME. (G) Comparison of the fitted parameter values for each marker (n= 20 cells/marker). (H) ROC curve generated from the binary logistic regression model built using the standard deviation, amplitude, and baseline. The closer the curve is to the upper left corner, the higher the predictive power of the parameter. (I) Sensitivity and specificity curves for a range of polarization cut-off values of SD. (J) Result of the classification of the training data set using a SD cut-off value of 60°. The region classified as polarized is marked in shaded orange. Scale bars shown in (B) and (F) are 5 μm. The following figure supplements are available for figure 1: **Figure supplement 1.** Localization pattern of polarity proteins used to test POME in *Arabidopsis* stomatal lineage. **Figure supplement 2.** Additional polarity parameters and classification models.

The macro calculates an angle of polarization with respect to the field of view after extracting the centroid and center of mass of a cell from the outline marker and the cortical polar protein channels, respectively. A line from the centroid to the center of mass defines the 0° and 180° angles used for reporting results (Figure 1D). Angles of polarization can be further analyzed to determine whether there is tissue-level coordination (Bringmann & Bergmann, 2017; Mansfield et al., 2018).

Initially, we used POME to quantify the distribution of BRXL2 in stomatal lineage ground cells (SLGCs) and found that its protein abundance at the membrane is well represented by a Gaussian model (Figure 1E). This allows us to describe the BRXL2 localization with three parameters: standard deviation (SD, *σ*), amplitude (*α*), and baseline intensity (*β*) (Figure 1E).

To evaluate the capacity of POME to distinguish among degrees of polarity, we applied it to three additional cases: BASL (polarized), BRXL2 during guard mother cell (GMC) divisions (depolarized), and myristoylated BRX (Myr-BRX, depolarized) (Figure 1F, Figure 1—figure supplement 1). These cases present a variety of challenges to polarity quantification, including signal interference from the nucleus (BASL), weak expression causing a low signal-to-noise ratio (depolarized BRXL2 in GMC), and signals from abutting membranes in two neighboring cells interfering with each other (myr-BRX). We selected 20 cells for each marker and measured their distribution along the cell periphery. The distributions of these markers were different, but they can all be described by a Gaussian model (Figure 1F). The SD, amplitude, and baseline intensity (Figure 1G) for BASL and the polarized BRXL2 were statistically indistinguishably (Figure 1—figure supplement 2A). The distribution of the depolarized (cytosolic) BRXL2 and Myr-BRX were clearly different from each other and from the distribution of the polarized BASL and BRXL2. Notice that the SD of depolarized BRXL2 and Myr-BRX are statistically indistinguishable, but their amplitudes and baselines are not, reflecting differences in fluorescence levels and possibly protein abundance (Figure 1G).

Comparing the SD, amplitude, and baseline intensity between our polarized and depolarized datasets using binary logistic regression suggest that SD alone is predictive of polarity (Figure 1H-I). Furthermore, the SD is nearly independent of the expression level of the marker and the imaging setting. A cut-off of 42° for SD can correctly discriminate between a polarized and a depolarized marker with a 95% classification sensitivity (true positive rate) and 83% specificity (true negative rate), while a cut-off of 60° for SD achieved 100% classification sensitivity and 78% specificity (Figure 1I, J). Alternatively, a support vector machine (SVM) algorithm can be used to generate a polarity classification model with 100% classification sensitivity and 100% specificity using the SD and normalized amplitude (*α*’, amplitude divided by mean fluorescence intensity) (Figure 1—figure supplement 3B-C).

### Characterizing the dynamics of polarity proteins in the stomatal lineage

We implemented POME to analyze the dynamics of BASL and BRXf distribution. Biochemical studies indicate that BASL and BRXf interact, while genetic analysis indicates that they require each other to remain polarized (Rowe et al., 2019). BASL and BRXf have been shown to occupy the same general region of the cortex, but the evidence is limited to single time point images and qualitative interpretation. As a result, it is unclear if their polarity always overlap or whether one protein polarizes first and then recruits the other.

To address this question, we first measured the distribution of BRXL2 before and after an ACD, from the initial formation of its polar crescent to its complete dissociation from the membrane (Figure 2A). We fitted the fluorescence measurements of each time frame and extracted the SD, amplitude, and baseline (Figure 2B). BRXL2 became polarized approximately 5±2.5 hours before the formation of the new cell wall and remained polarized for over 9±2.5 hours afterward, using the SD cut-off of 60° from our binary logistic regression. The SD and baseline values of the BRXL2 distribution remained remarkably constant over this period, while the mean amplitude increased and then decreased monotonically. This result could be consistent with scenarios where BRXL2 is locally delivered to the crescent and retained there, or uniformly delivered to the membrane but transported into the crescent more rapidly than it can diffuse away.

**Figure 2.**
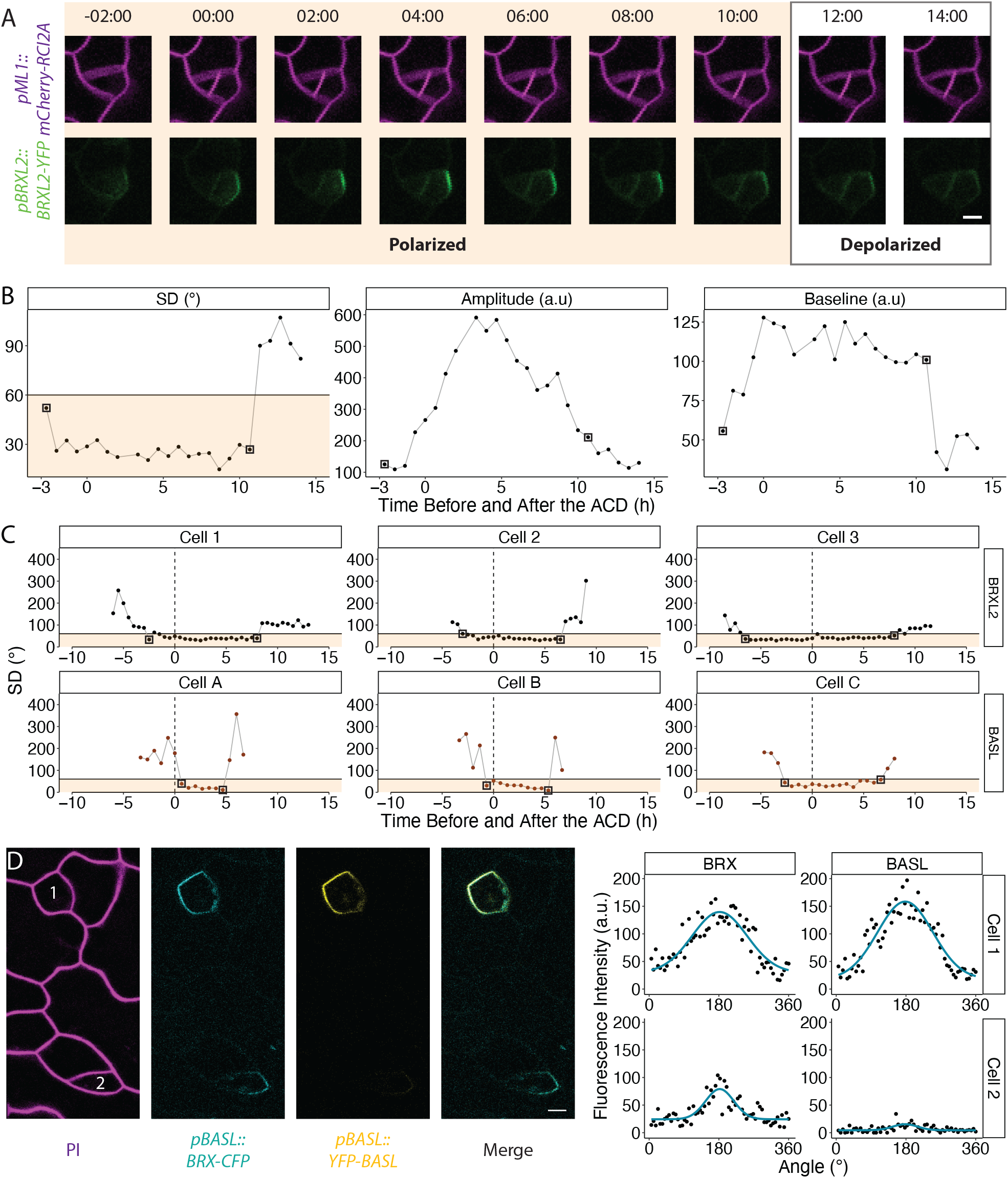
POME reveals differences in BRXL2 and BASL polarity dynamics during ACDs. (A) Time course of BRXL2 localization during an ACD. Cells are dually marked by *pML1::RCI2A-mCherry* (magenta) and *pBRXL2::BRXL2-YFP* (green). Time point 00:00 (hours:minutes) marks the formation of the cell plate. (B) Changes in SD, amplitude, and baseline of the BRXL2 distribution determined by POME. (C) Plotted measurements of individual cells, each expressing either BRXL2 (top cells, 1-3) or BASL (bottom cells, A-C). Vertical dashed lines indicate cell plate formation. Solid horizontal lines indicate a SD cut-off value of 60°, and all values in shaded orange are classified as polarized. The 60° SD cut-off value that achieved 100% classification sensitivity is chosen because of the lower image resolution and lower signal to noise ratio of time-lapse images compared to still images. For each cell, the first and last time-points showing polarization are marked by black boxes. (D) BRX (cyan) and BASL (yellow) localization pattern during the late G2/early mitotic phase in two meristemoids. Note that in cell 2, BRX is polarized while BASL is not. Scale bars shown in (A) and (D) are 5 μm. The following figure supplement is available for figure 2: **Figure supplement 1.** Confocal images of BRXL2 and BASL localization pattern during ACDs.

Subsequently, we performed a similar characterization of BASL dynamics. Comparison of the dynamics of BRXL2 and BASL showed that BRXL2 polarized earlier than BASL during ACDs, and stayed polarized longer than BASL afterward (Figure 2C, Figure 2—figure supplement 1). A dual-reporter line expressing BRX-CFP (a homolog of BRXL2 redundant in the stomata lineage) and BASL-YFP further supports the hypothesis that BRXf polarizes ahead of BASL during ACDs. In still images of the dual-reporter line, from which we could obtain higher resolution images than in time-lapse, we identified a few cells at late G2/ early mitotic phase where BRX, but not BASL, was polarized (Figure 2D).

We also examined the dynamics of BRXL2 in *basl-2* null mutants, where BRXL2 was believed to be depolarized (Rowe et al., 2019). Strikingly, we found that in some *basl-2* cells, BRXL2 could still polarize transiently during ACDs (Figure 3). Whether this transient BRXL2 polar crescent in *basl-2* is functional, however, is debatable, since it does not show up in all the dividing cells and the *basl-2* mutant has evident ACD defects (Dong et al., 2009). Nevertheless, these results suggest that BRXL2 accumulates at the polar crescent first and then recruits BASL, and the formation of their protein complex is required to retain both proteins at the crescent.

**Figure 3.**
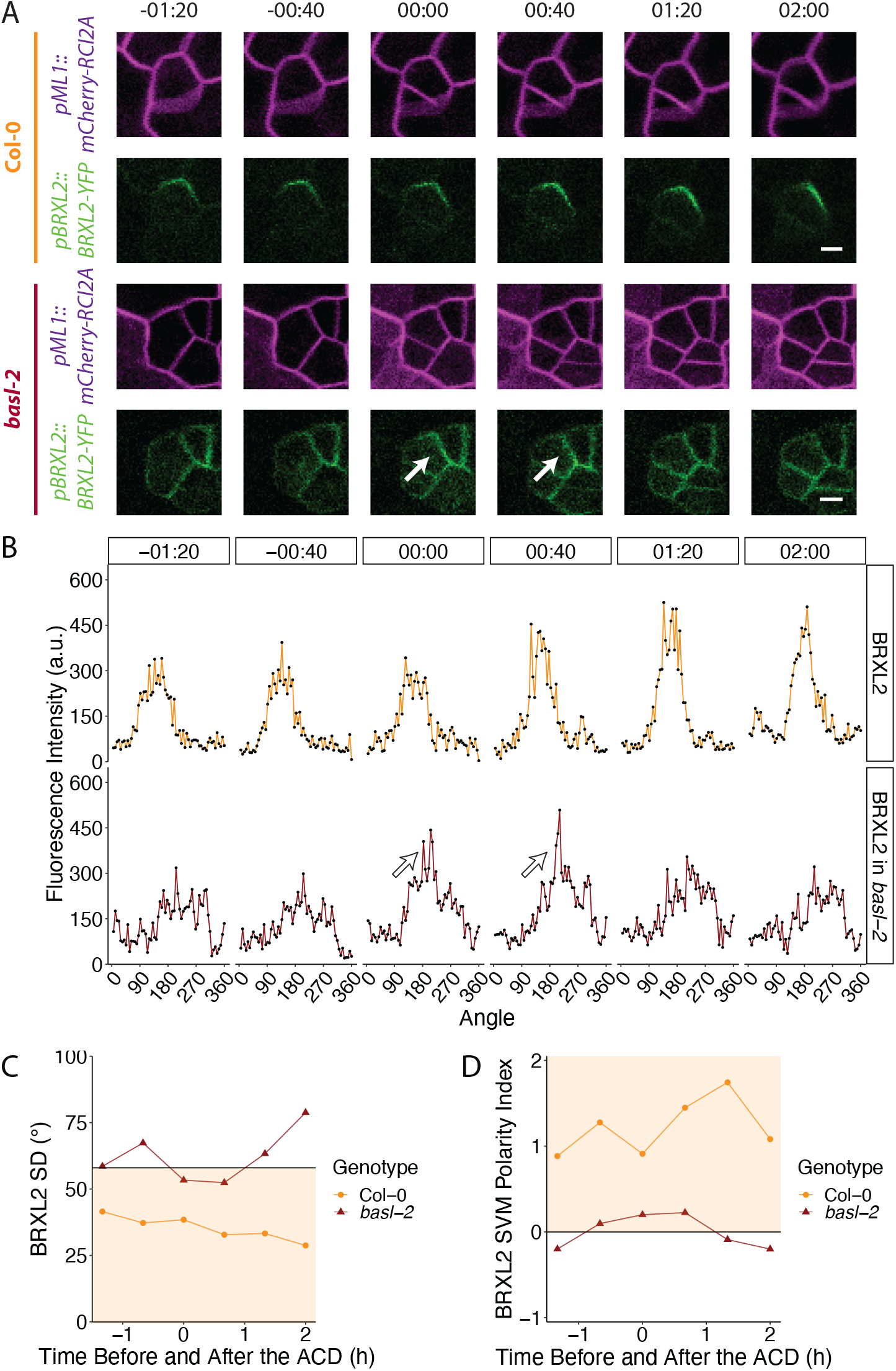
POME shows BRXL2 can transiently polarize during ACDs in basl-2 mutant. (A) BRXL2 localization pattern during cell division in wild-type (Col-0) and the *basl-2* mutant. Cells dually marked by *pML1::RCI2A-mCherry* (magenta) and *pBRXL2::BRXL2-YFP* (green) are tracked before, during and after ACD, where 00:00 (hours: minutes) corresponds to the formation of the cell plate. (B) POME measurements of BRXL2 polarity from cells in (A). The formation of a transient BRXL2 polar crescent in *basl-2* at time frame 00:00 and 00:40 is indicated by a white arrow in (A) and (B). (C-D) BRXL2 SD and SVM polarity index estimated for each time point in (A). The decision boundary of each classification model is labeled with solid black lines, and the region classified as polarized is marked in shaded orange. Scale bar in (A) is 5 μm.

### Applications of POME in other developmental contexts

POME was initially designed to address the challenges of measuring polarity in dispersed, asynchronously dividing, leaf epidermal cells. An automatic tool to quantify polarity, however, is useful for other developmental contexts, and the simple design of POME should make it adaptable to other tissues and organisms.

In *Arabidopsis*, the MAPKKK YODA (AtYDA) is polarized and segregated preferentially to larger daughter cells in the stomatal lineage ACDs by BASL (Zhang et al., 2015). In *Brachypodium*, a YODA homolog, BdYDA1, is also required for stomatal lineage ACDs (Abrash et al., 2018). However, the *Brachypodium* genome does not encode a BASL homolog (Bowles et al., 2020). It was therefore unclear whether BdYDA1 would be polarized in the *Brachypodium* stomatal lineage. Grass stomata are arranged in cell files and exhibit a base-to-tip gradient of development. Meristemoids undergo directional ACDs such that the smaller cell (stomatal precursor) is oriented towards the tip and the larger one (pavement cell precursor) towards the base (Fig 4A-B). To determine if BdYDA1 is polarized, we used POME to quantify its distribution in individual meristemoids before ACDs and in the newly formed larger daughter cells resulting from this division (Figure 4A-B).

**Figure 4.**
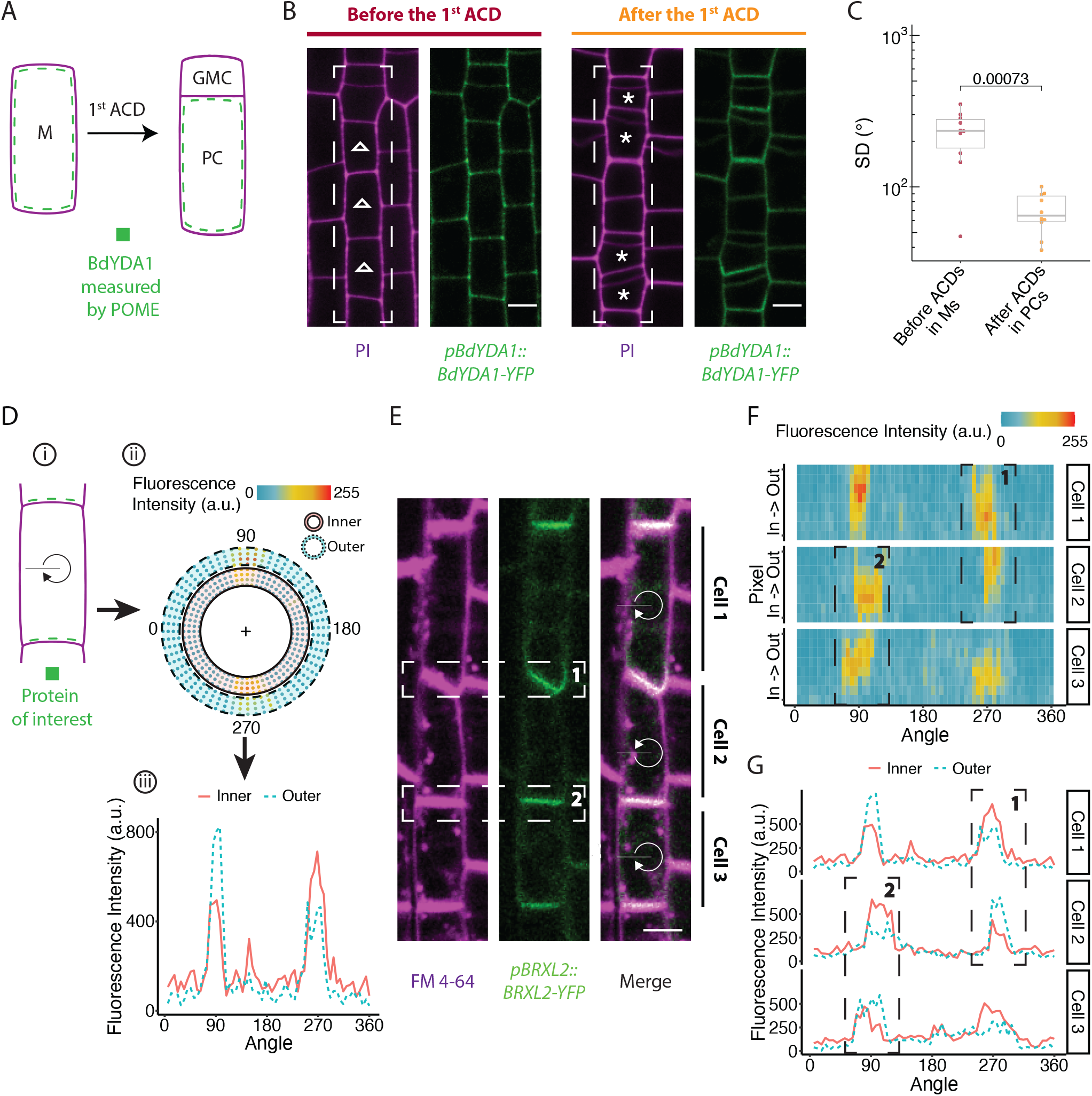
Application of POME to quantifying polarity in diverse cells. (A-C) POME reveals BdYDA1 is polarized in *Brachypodium* stomatal lineage cells. (A) Schematic of the cell types in the *Brachypodium* stomatal lineage. Meristemoids (M) divide asymmetrically to generate one small guard mother cell precursor (GMC) and one large pavement cell precursor (PC). (B) *pBdYDA1::BdYDA1-YFP* (green) localization pattern in stomatal lineage cells (indicated by white-dashed boxes) before (left) and after (right) the first ACD. Cell boundaries are visualized by propidium iodide (PI, magenta). Ms and PCs selected for POME analysis are labelled with open triangles (Ms) and asterisks (PCs). (C) Plot of the SD indicates that BdYDA1 is not polarized in meristemoids (pre-ACD) but is polarized in pavement cell precursors. (n=10 cells/stage). (D-G) Application of POME to determine shootward or rootward polarity in contiguous cells. (D) Scheme of how POME can be used to determine which of two adjacent cells is expressing a polarized marker. i) A target cell shows localization of the polarized marker (green) both within its membrane (below) and adjacent to its membrane (above). The black line indicates the 0° angle and the clockwise arrow indicates the direction of rotation for POME measurement. ii) Reconstruction of marker localization in inner and outer membrane regions. Fluorescence in the inner membrane region (within the solid rings) is contributed by the selected cell; while fluorescence in the outer membrane region (within the dashed rings) is considered to be contributed by the neighboring cell. iii) Comparison of the marker intensity (sum of all pixel values) in the inner (solid red line) and outer regions (dashed blue line) at a particular angle indicate whether the signal at the membrane is mostly coming from the neighboring cell (as shown at 90°) or from the target cell (as shown at 270°). (E) Rootward localization of BRXL2 (green) in three adjacent lateral root cap cells in *Arabidopsis*. Plasma membrane visualized with FM4-64 (magenta). Regions of interest (ROIs) analyzed to determine on which side of the shared interphase a polarized protein resides are indicated by dashed boxes labeled as 1 and 2 in (E), (F), and (G). (F) Heat map of the fluorescence intensity of the BRXL2 localization shown in (E). For each angle, the four innermost and outermost pixels are considered the inner and outer membrane regions, respectively. (G) For the three cells shown in (E), the total fluorescence intensities of the inner (solid red line) and outer (dashed blue line) membrane regions are plotted at each angle. Scale bars in (B) and (E) are 5 μm. The following figure supplements are available for figure 4: **Figure supplement 1.** Application of POME in root epidermal cells to validate shootward localization of PIN2. **Figure supplement 2.** Application of POME is quantifying PAR-2 polarity in the *C. elegans* P cell lineage during embryo development.

BdYDA1 was not polarized in meristemoids, but after ACD, polarized BdYDA1 was detected in the larger daughter cell (Fig 4B-C). This segregation and polarization of BdYDA1 is consistent with a role restricting stomatal fate, and these larger daughters often wrongfully acquire stomatal identity in the loss-of-function mutant *bdyda1* (Abrash et al., 2018). The position of polar BdYDA1, distal to the new division plane in the larger daughter cell, is equivalent to where AtYDA localizes in a BASL-dependent manner in *Arabidopsis* (Zhang et al., 2015). This leads to the interesting conclusion that MAPKKKs can and must polarize during stomatal ACDs to ensure differential fates in different plants, but BASL may be newly recruited into this role in dicots and alternative mechanisms must exist to polarize BdYDA1.

The expression of BdYDA1 in each of many contiguous stomatal cells represented a complication for automated polarity analysis. We had to rely on prior knowledge about oriented ACDs (Abrash et al., 2018) to infer in which cells and on what cell face BdYDA1 was localized. The problem of assigning polarity to individual contiguous cells is quite common. In *Arabidopsis* roots, for example, cells are also arranged in files, and localization of proteins to specific faces is required for the development and physiological functions of the root. Many proteins in the root and vascular systems, especially those involved in polar auxin transport, display polar localization to “shootward” and/or “rootward” faces of the cell (Breda et al., 2017; Marhava et al., 2018; Scacchi et al., 2009; Wisniewska et al., 2006; Zourelidou et al., 2009) (Figure 4E, Figure 4—figure supplement 1A). Determining whether a protein is present at both shootward and rootward faces in a cell within a contiguous file has been challenging.

To test whether POME could be used to determine which side of the interphase between two adjacent cells a protein resides, we quantified the localization of BRXL2 and the transmembrane protein PIN2 in *Arabidopsis* roots. BRXL2 has been reported to localize rootward in roots (Marhava et al., 2020) and PIN2 to localize shootward in root epidermis cells (Wisniewska et al., 2006). We quantified the pixel intensity at multiple “rings” surrounding the plasma membrane to ascertain whether a polarized protein is enriched in either outer or inner rings of a particular cell. With this modified POME protocol, we validated BRXL2’s rootward localization pattern in lateral root cap cells (Figure 4E-G) and PIN2’s shootward localization in root epidermal cells (Figure 4—figure supplement 1). The same principle of capturing pixel intensity of a reporter across the membrane could also be used to determine the redistribution of proteins between membrane and cytoplasm, a common response to environmental perturbation in plants (Marhava et al., 2018; Zourelidou et al., 2009).

## CONCLUSION

Plant proteins exhibit staggering variation in cortical polar domains. They can be found linked to dispersed ACDs, in contiguous cell files aligned with organ axes, or nestled in cell corners. POME can quantify dynamic polarity in these different contexts and complements other recent advances in cell segmentation and quantitative image analysis (Wolny et al., 2020). POME converts a once low-throughput and qualitative procedure into a semi-automated one that is amicable to statistical analysis. It could be even more powerful when combined with other techniques such as FRAP and FRET-FLIM to test models of polarity formation.

Ultimately, we expect fluorescence microscopy and tools like POME to answer some and open new questions in cell biology. In our exploration of stomatal lineage proteins in two plant species, we were able to find that BASL, while a central hub in stomatal polarity, may be preceded by BRXL2 polarity in *Arabidopsis* and must be supplanted by other mechanisms for establishing polar localization of the MAPKKK BdYDA1 in *Brachypodium*. POME also allowed us to validate the rootward and shootward localization of BRXL2 and PIN2, respectively, in *Arabidopsis* roots. Furthermore, the utility of POME is not limited to plants, as we were also able to use it to quantify PAR-2 polarity in *C. elegans* embryos (Hubatsch et al., 2019) (Figure 4—figure supplement 2). The simplicity and versatility of our pipeline make it accessible to the broad research community regardless of previous programming skills.

## MATERIALS AND METHODS

### Key resources table

| Reagent | Designation | Source or Reference | Identifiers | Additional Information |
| --- | --- | --- | --- | --- |
| Gene (*Arabidopsis thaliana*) | BASL | NA | AT5G60880 | Polarity protein in the stomatal lineage |
| Gene (*Arabidopsis thaliana*) | BRXL2 | NA | AT3G14000 | Polarity protein in the stomatal lineage |
| Gene (*Arabidopsis thaliana*) | BRX | NA | AT1G31880 | Polarity protein in the stomatal lineage and the phloem |
| Gene (*Arabidopsis thaliana*) | ML1 | NA | AT4G21750 | Homeobox transcription factor |
| Gene (*Arabidopsis thaliana*) | PIN2 | NA | AT5G57090 | Auxin efflux carrier |
| Gene (*Brachypodium distachyon*) | BdYDA1 | NA | Bradi5g18180 | MAPKKK |
| Gene (*Caenorhabditis elegans*) | PAR-2 | NA | CELE_F58B6.3 | Polarity protein; WormBase ID: WBGene00003917 |
| Strain, strain background<br>(*Arabidopsis thaliana*) | Col-0 | ABRC | CS28166 | Wild-type *Arabidopsis* ecotype used in this study |
| Strain, strain background<br>(*Agrobacterium tumefaciens*) | GV3101 | other |  | Electrocompetent *A. tumefaciens* |
| Genetic reagent (*Arabidopsis thaliana*) | *basl-2* | Dong et al., 2009; DOI: 10.1016/j.cell.2009.04.08 | Wisc sLox264F02 |  |
| Genetic reagent (*Arabidopsis thaliana*) | *pBRXL2::BRXL2-YFP; pML1::mCherry-RCI2A* | Rowe et al., 2009; DOI: 10.1101/614636 |  |  |
| Genetic reagent (*Arabidopsis thaliana*) | *pBASL::YFP-BASL; pML1::mCherry-RCI2A* | Rowe et al., 2019; DOI: 10.1101/614636 |  |  |
| Genetic reagent (*Arabidopsis thaliana*) | *pBASL::Myr-BRX-YFP; pML1::mCherry-RCI2A* | Rowe et al., 2019; DOI: 10.1101/614636 |  |  |
| Genetic reagent (*Arabidopsis thaliana*) | *pBRXL2::BRXL2-YFP; pML1::mCherry-RCI2A; basl-2* | Rowe et al., 2019; DOI: 10.1101/614636 |  |  |
| Genetic reagent (*Arabidopsis thaliana*) | *pBRXL2::BRXL2-CFP; pBASL-YFP-BASL; brxq* | Rowe et al., 2019; DOI: 10.1101/614636 |  |  |
| Genetic reagent (*Arabidopsis thaliana*) | *pPIN2::PIN2-GFP* | Wisniewska et al., 2006; DOI: 10.1126/science.1121356 |  |  |
| Genetic reagent (*Brachypodium distachyon*) | *pBdYDA1::BdYDA1-YFP* | Abrash et al., 2018; DOI:10.1242/dev.165860 |  |  |
| Dye | FM 4-64 | Thermo Fisher | Thermo Fisher: T3166 |  |
| Dye | Propidium Iodide | Thermo Fisher | Thermo Fisher: P3566 |  |
| Software, algorithm | Leica Application Suite X | Leica |  | Version: 3.5.2.18963 |
| Software, algorithm | FIJI | Schindelin et al., 2012; DOI: 10.1038/nmeth.2019 |  | Version: 2.0.0-rc-69/1.52p |
| Software, algorithm | POME | This manuscript |  | Version: 1.0.0 |
| Software, algorithm | R | https://www.R-project.org/ |  | Version: 4.0.1 |
| Software, algorithm | R studio | https://rstudio.com |  | Version: 1.3.959 |

### Plant material

All *Arabidopsis* lines used in this study were generated in the Col-0 background. Construction of all polarity protein reporter lines (*pBRXL2::BRXL2-YFP pML1::mCherry-RCI2A, pBASL::YFP-BASL pML1::mCherry-RCI2A, pBASL::Myr-BRX-YFP pML1::mCherry-RCI2A brxq, pBRXL2::BRXL2-YFP pML1::mCherry-RCI2A basl-2, pPIN2::PIN2-GFP*) have been reported elsewhere (Rowe et al., 2019; Wisniewska et al., 2006). The *Brachypodium* BdYDA1 reporter line *pBdYDA1::BdYDA1-YFP* was generated in the Bd21-3 ecotype as described elsewhere (Abrash et al., 2018).

### Plant growth conditions

All *Arabidopsis* seeds were surface sterilized by bleach or 75% ethanol and stratified for 2 days. After stratification, seedlings were vertically grown on ½ Murashige and Skoog (MS) media for 3-6 days under long-day condition (16h light/8h dark at 22 °C).

### Microscopy and image acquisition

All imaging experiments of *Arabidopsis* were performed on a Leica SP5 confocal microscope with HyD detectors using 25x NA0.95 and 40x NA1.1 water objectives with image size 1024*1024 and digital zoom from 1x to 2.5x. For time-lapse experiments, 3-day post germination (dpg) seedlings were mounted in a custom imaging chamber filled with ½ MS solution (Bringmann & Bergmann, 2017; Davies & Bergmann, 2014; Simmons et al., 2019). Laser settings for each reporter, except the membrane marker (*pML1::mCherry-RCI2A* or PI), were adjusted to avoid over-saturation. For the time-lapse experiments reported in this study, 30 min or 40 min intervals between each image stack capture were used.

Images of the BdYDA1 reporter in developing *Brachypodium* leaves were reanalyzed from the same set of raw images as reported in Figure 3 of Abrash et al., 2018. Images of PAR dynamics in *C. elegans* embryos were reanalyzed from the raw images of Supplementary Video 2 reported in Hubatsch et al., 2019.

### POME analysis of fluorescence images

All raw fluorescence image Z-stacks were projected with Sum Slices in FIJI unless noted otherwise. For all time-lapse images, drift was corrected using the Correct 3D Drift plugin (Parslow et al., 2014). These time-lapse images were then split into individual time frames and treated as still images for analysis afterwards. The channels of images for POME analysis were reorganized, where the cell outline marker channel was placed in channel one and the polarity protein channel was placed in channel two. Each individual cell of interest was then examined for manual correction before passing to POME, when regions containing interfering vesicles (images with FM4-64 staining) or nucleus (images of BASL reporter) were manually removed. When running POME on a selected cell, the settings were adjusted per experiment. Information on how to determine POME settings is described in detail in the Appendix 1. The results of POME measurement were then imported, summarized, and analyzed in RStudio.

Fitting the polarity protein distribution along the cell membrane to a Gaussian model For polarity proteins in *Arabidopsis* and *Brachypodium* stomatal lineage, the mean florescence intensity of the polarity protein reporter along the cell membrane was fitted to a Gaussian model (Equation 1) by non-linear regression with nls function from the stats package (R Core Team, 2020). Key parameters of this regression model, including standard deviation *σ*, center *μ*, amplitude *α*, and baseline value *β*, were estimated using the formula:

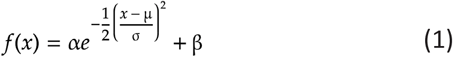

### Generating polarity classification matrixes

To determine if the parameters (*σ*, *α*, and *β*) estimated by POME were sufficient to distinguish between polarized (BRXL2 in SLGCs and BASL) and depolarized (BRXL2 during GMC divisions and Myr-BRX) markers, we used measurements from 80 cells (20 cells per marker) to train a binary logistic regression model using the glm function from the stats package in RStudio. To visualize each parameter’s ability to classify polarity, a receiver operating characteristic (ROC) plot was generated with the roc function from the pROC package (Robin et al., 2011). The resulting plot (Figure 1H) suggest that *σ* alone can successfully discriminate between polarized and depolarized markers. Next, a simpler binary logistic regression model was built using only *σ* (Figure 1I) to determine a suitable cut-off value. To calculate the error rate of the polarity *~ σ* logistic regression model, a 10-fold cross-validation prediction error was calculated with the training dataset with the cv.glm function in the boot package (Davison & Hinkley, 1997),where an average error rate of 9.2% prediction error was reported.

For the support vector machine (SVM) model, the parameters from the same 80 cells in the binary logistic regression model were used as training data. To remove imaging setting and reporter basal expression bias, a normalized amplitude *α*’ was calculated by dividing the amplitude α by the mean fluorescence intensity of all angles. The SVM model was then built with the svm function in the e1071 package (Meyer et al., 2019). Next, linear coefficients were extracted from the model to calculate the slope (3.672) and intercept (0.069) of the decision boundary (Figure 1—figure supplement 2B). We can then define an SVM polarity index as the distance to the SVM model decision boundary calculated with the formula:

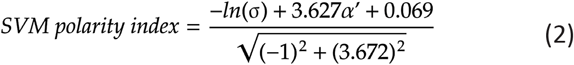

A positive SVM polarity index indicates a polarized cell, while a negative a SMV polarity index indicates a depolarized cell. Higher SMV polarity index values indicate higher polarization.

### Statistical analysis

All statistical analyses were performed in RStudio. Unpaired Mann-Whitney tests were conducted to compare two data samples with the compare_means function from the ggpubr package (Kassambara, 2020).

## Acknowledgments

We thank Dr. Nathan Goehring for providing the raw images of PAR protein behaviors in *Caenorhabditis elegans* embryo and Ximena Anleu-Gil for images of BdYODA1 in *Brachypodium*. We thank Katelyn McKown, Gabriel Amador, Dr. Andrea Mair, and other members of the Bergmann lab for valuable feedback on the manuscript. This material is based upon work supported by the National Science Foundation under Grant No. 1942722 to Lily S. Cheung.

## ADDITIONAL INFORMATION

### Funding

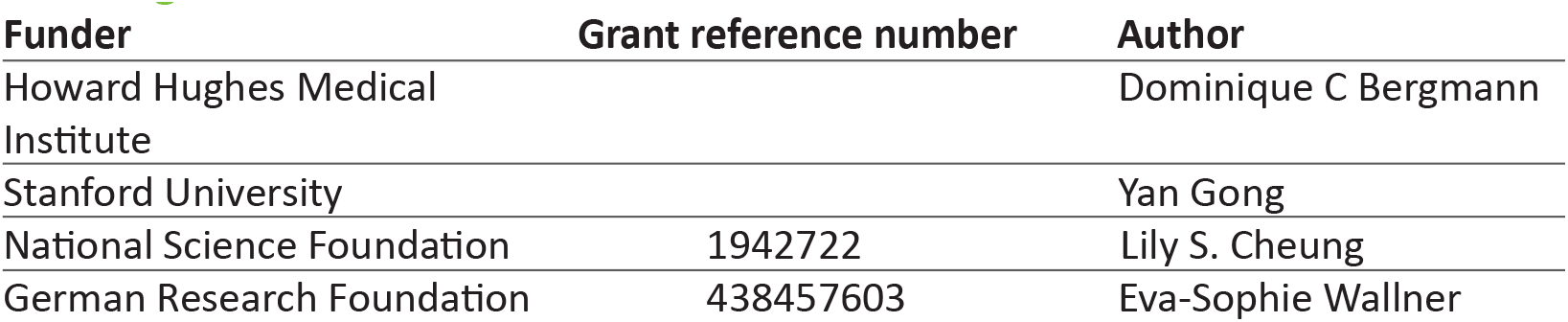

### Author contributions

Yan Gong, Conceptualization, Data curation, Formal analysis, Validation, Visualization, Methodology, Writing—original draft, Biological experiments and imaging, Design of image quantification pipeline, Image analysis, Interpretation of results; Rachel Varnau, Data curation, Formal analysis, Biological experiments and imaging, Image analysis, Interpretation of results; Eva-Sophie Wallner, Conceptualization, Data curation, Formal analysis, Biological experiments and imaging, Image analysis, Interpretation of results; Dominique C. Bergmann, Supervision, Funding acquisition, Project administration, Conceptualization, Data curation, Formal analysis, Writing—original draft, Design of image quantification pipeline, Interpretation of results; Lily S. Cheung, Supervision, Funding acquisition, Project administration, Conceptualization, Data curation, Formal analysis, Methodology, Writing—original draft, Design of image quantification pipeline, Image analysis, Interpretation of results.

---

**Figure 1—figure supplement 1.**
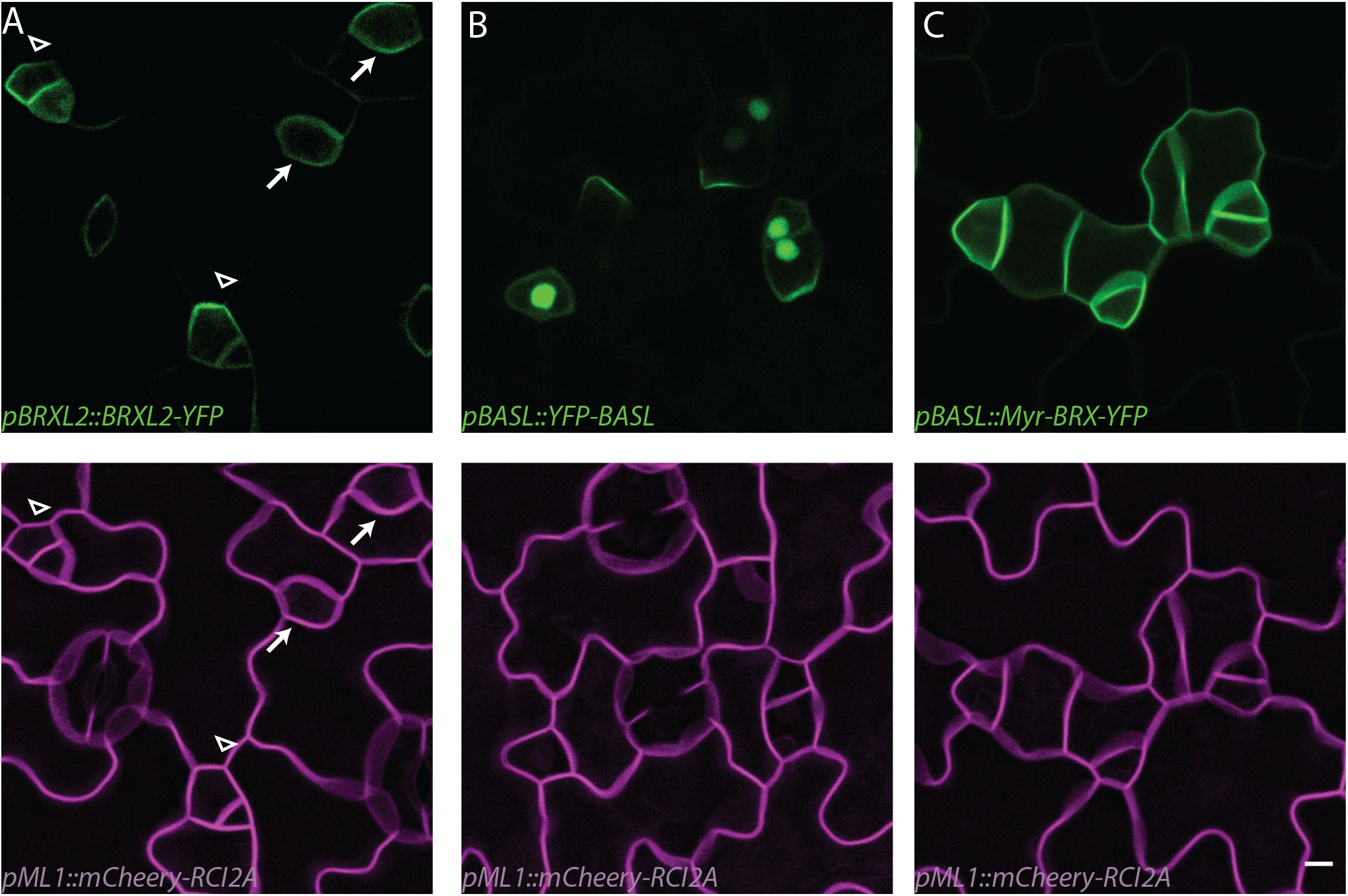
Localization pattern of polarity proteins used to test POME in Arabidopsis stomatal lineage. (A-C) Representative confocal images showing larger areas than Figure 1F. Fluorescent markers used to test POME. Magenta: *pML1::RCI2A-mCherry*; green: *pBRXL2::BRXL2-YFP* (A), *pBASL::YFP-BASL* (B), *pBASL::Myr-BRX-YFP* (C). In (A), SLGCs with polarized BRXL2 are labeled with arrowheads and GMCs with depolarized BRXL2 are labeled with arrows. Scale bar is 5 μm.

**Figure 1—figure supplement 2.**
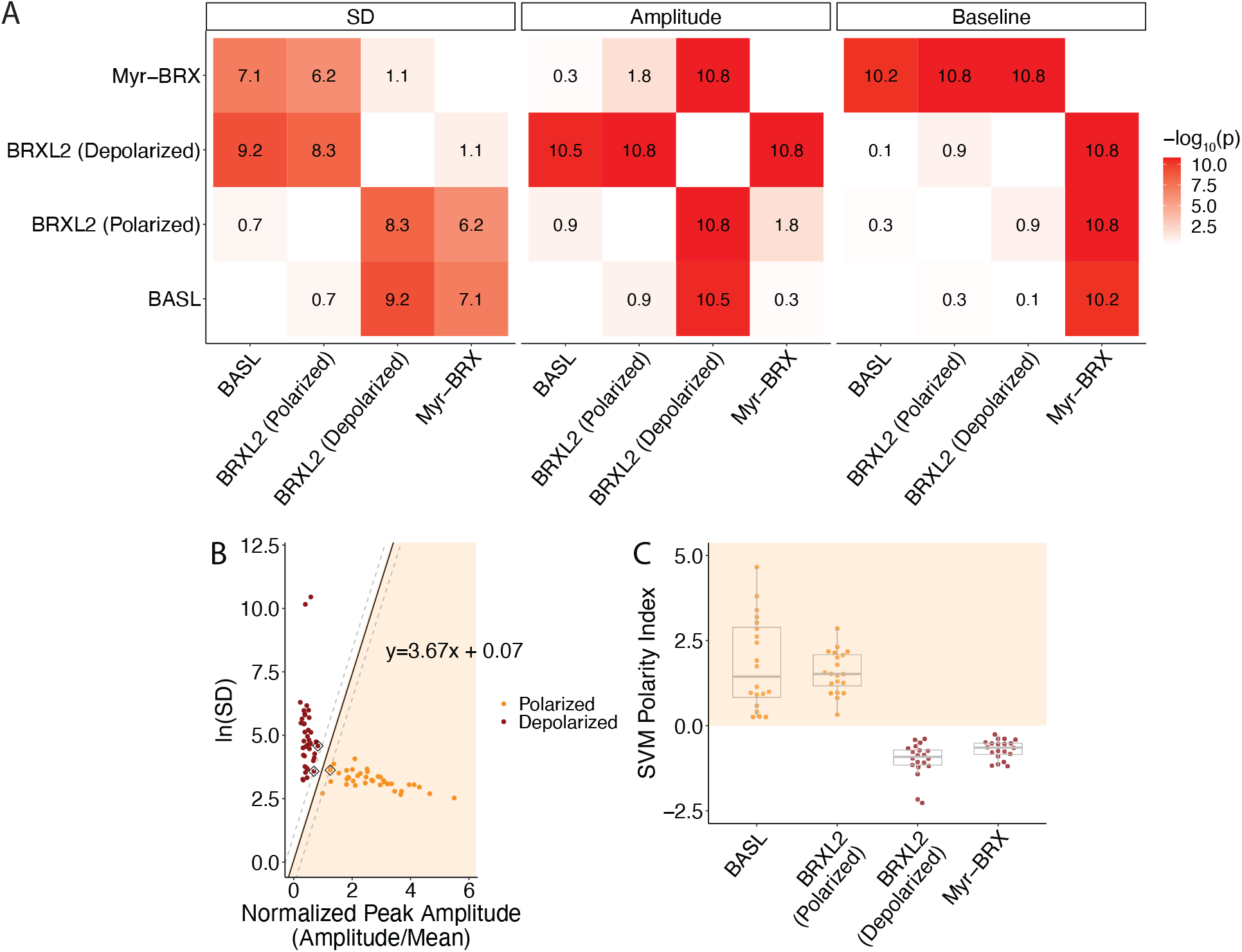
Additional polarity parameters and classification models. (A) p-values calculated by unpaired Mann-Whitney tests comparing each of the Gaussian model parameters across all the polarity markers shown in Figure 1F. (B) SVM classification model based on the SD (*σ*) and normalized amplitudes (*α*’) estimated for the cells in Figure 1G. Each cell is represented by a solid dot, the support vectors are marked by black squares, the decision boundary is labeled by the solid black line, the margin boundaries are labeled with gray dash lines, and the classified polarized region is marked in shaded orange. Deviation from the decision boundary line to the right indicates higher polarity. (C) SVM polarity index of the four polarity markers shown in Figure 1F. The classified polarized region is marked in shaded orange.

**Figure 2—figure supplement 1.**
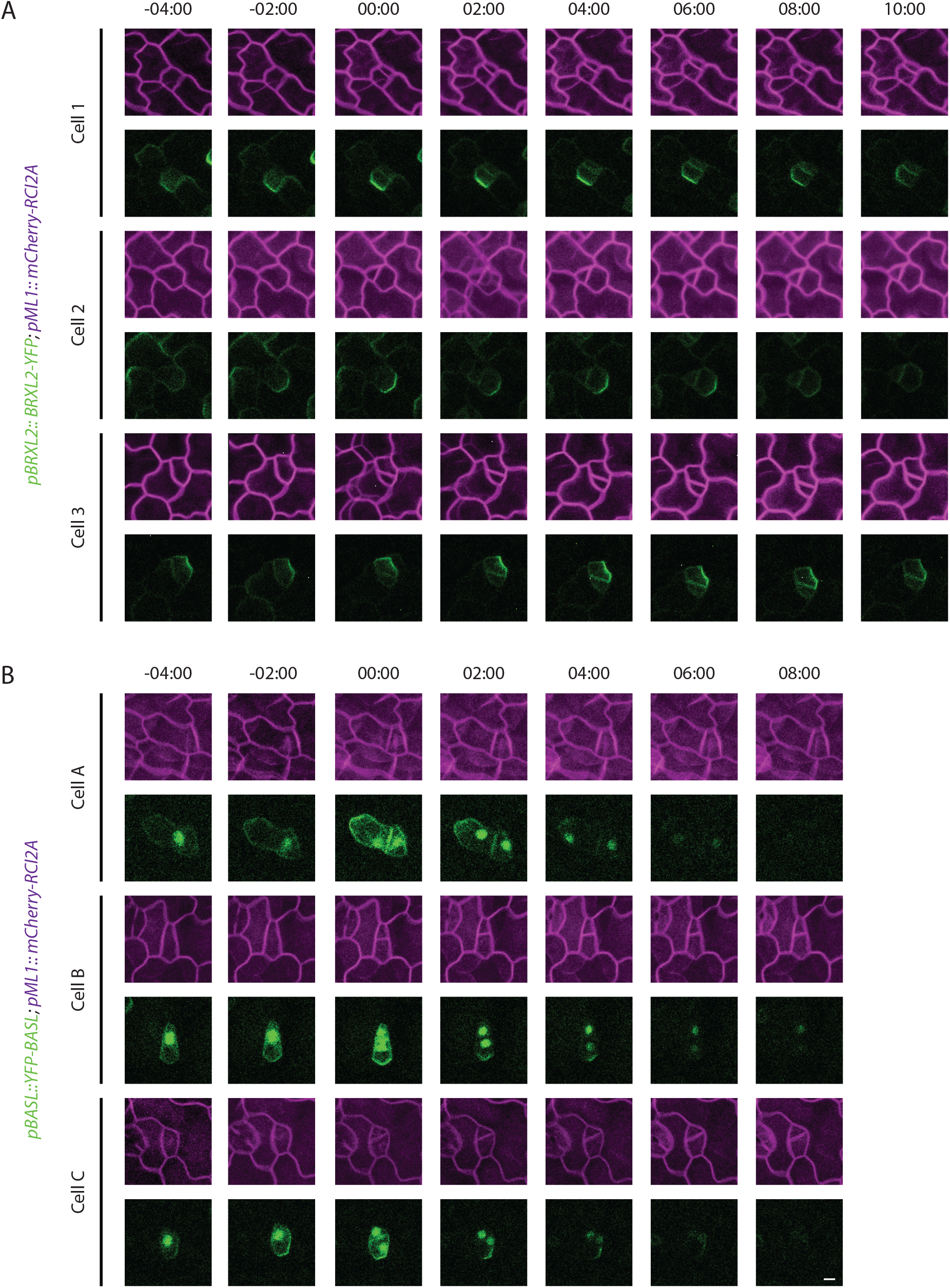
Confocal images of BRXL2 and BASL localization pattern during ACDs. (A-B) BRXL2 (A) and BASL (B) localization pattern during the six ACDs quantified in Figure 2C. Cells dually marked by *pML1::RCI2A-mCherry* (magenta) and *pBRXL2::BRXL2-YFP* (green, A) or *pBASL::YFP-BASL* (green, A) are tracked before, during and after ACDs. Timepoint 00:00 (hours: minutes) marks the formation of the cell plate. Scale bar is 5 μm.

**Figure 4—figure supplement 1.**
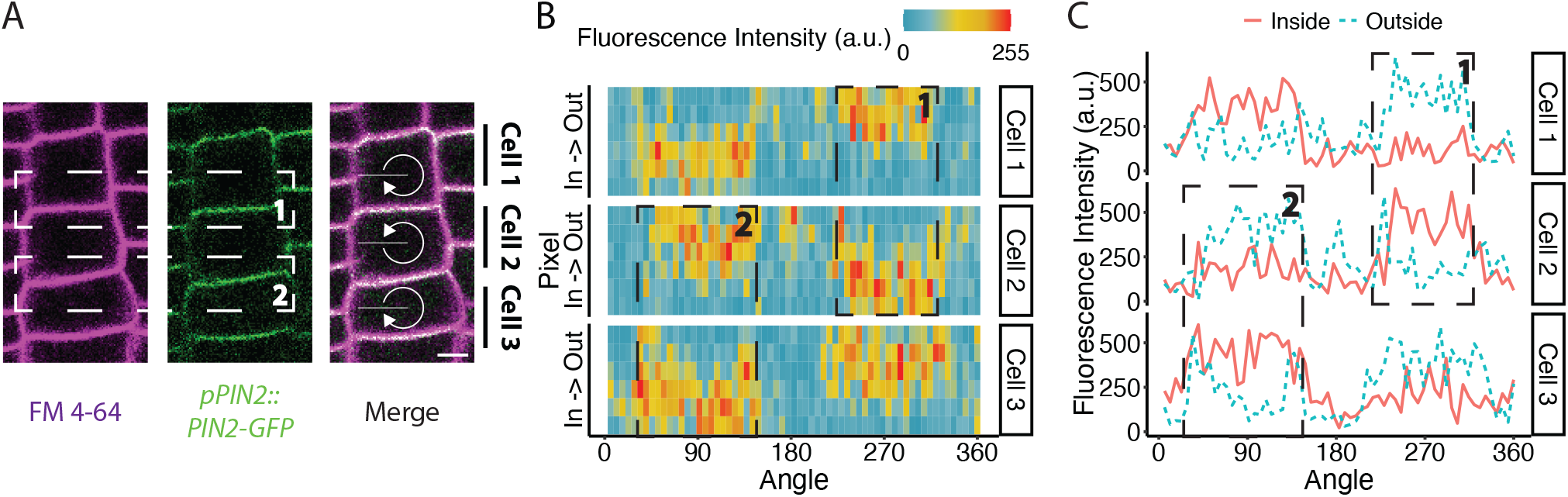
Application of POME in root epidermal cells to validate shootward localization of PIN2. (A) Localization of PIN2 (green) in three adjacent epidermal cells in Arabidopsis root epidermal cells. Plasma membrane visualized with FM4-64 (magenta). ROIs analyzed to determine on which side of the shared interphase a polarized protein resides are indicated by dashed boxes labeled 1 and 2 in all panels. (B) Heat map of the fluorescence intensity of the PIN2 localization shown in (A). For each angle, the three innermost and outermost pixels are considered the inner and out membrane regions, respectively. (C) For the three cells shown in (A), the total fluorescence intensities of the inner (solid red line) and outer (dashed blue line) membrane regions are plotted at each angle. Comparison of the total fluorescence intensity in the inner and outer membrane regions suggest that PIN2 is localized to the shootward face of cells. Scale bar in (A) is 5 μm.

**Figure 4—figure supplement 2.**
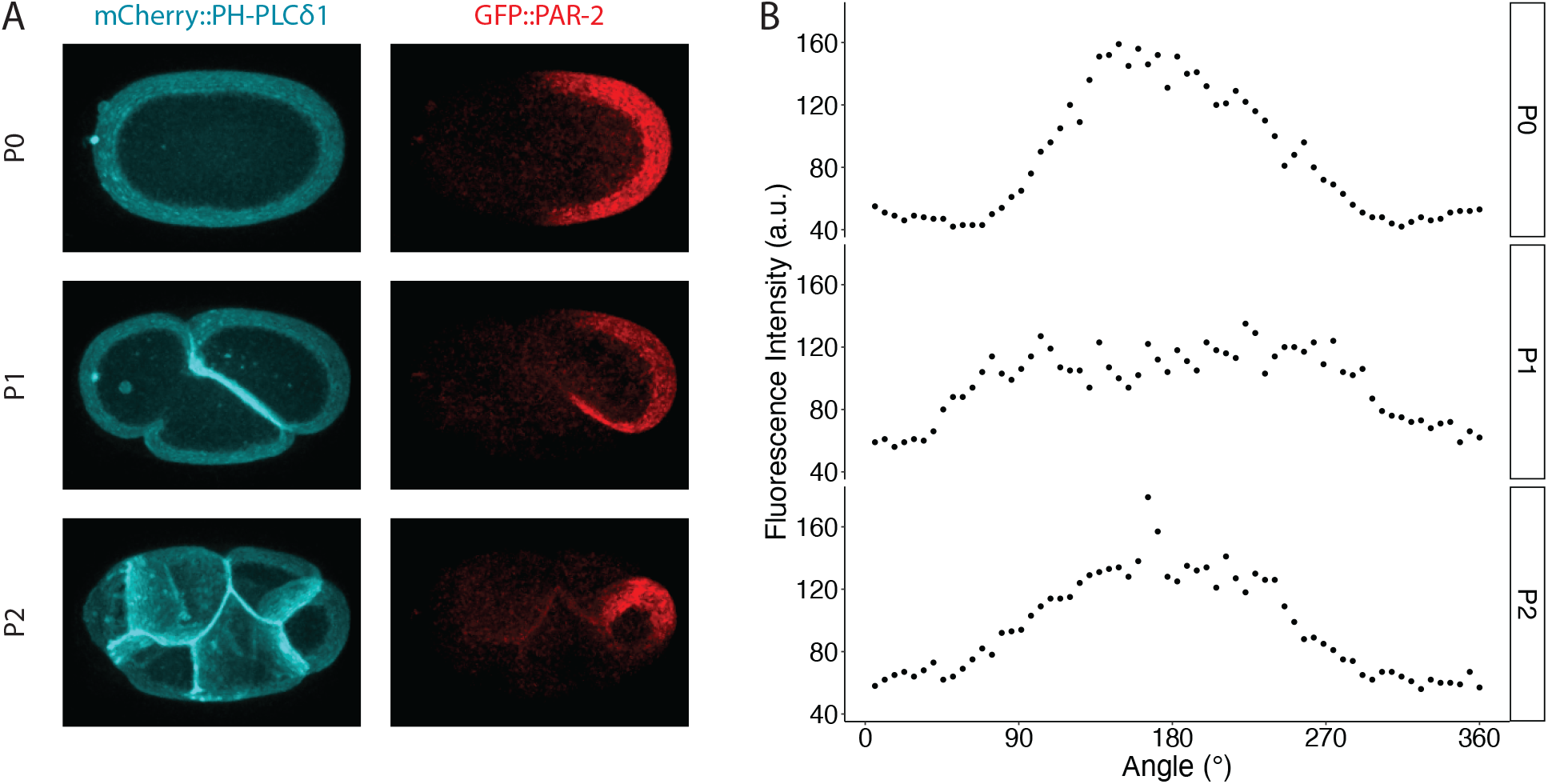
Application of POME is quantifying PAR-2 polarity in the *C. elegans* P cell lineage during embryo development. (A-B) Confocal images (A) and POME measurements (B) PAR-2 polarity in consecutive P0, P1, P2 cells during *C. elegans* embryo development. The confocal images in (A) correspond to three different frames from the Supplementary Video 2 accompanying Hubatsch et al., 2019.

